# Quorum-sensing gene regulates hormetic effects induced by sulfonamides in Comamonadaceae

**DOI:** 10.1101/2023.03.31.535187

**Authors:** Hui Lin, Xue Ning, Donglin Wang, Qiaojuan Wang, Yaohui Bai, Jiuhui Qu

## Abstract

Hormesis is a toxicological phenomenon whereby exposure to low-dose stress results in stimulation of various biological endpoints. Among these, the induction of cell proliferation by antibiotics is critical, but the underlying molecular mechanisms remain poorly understood. Here, we showed that sulfonyl-containing chemicals (e.g., sulfonamides) can induce cell-proliferation hormesis of *Comamonas testosteroni*. Investigation of the hormesis mechanism revealed that low-dose sulfonamides bind to the *LuxR*-type quorum sensing protein LuxR solo, thereby triggering the transcription of 3-ketoacyl-CoA thiolase, a key enzyme of the fatty acid β-oxidation. This provides additional ATP, NADPH, and acetyl-CoA for purine and pyrimidine biosynthesis, allowing cells to synthesize sufficient nucleotides to support rapid cell growth. Our work reports on a previously unknown mechanism for the hormetic effect and highlights its generality in the Comamonadaceae family.

## Introduction

Hormesis commonly refers to the adaptive response of organisms (e.g., microbial, plant, and animal species) to low-dose stressors (e.g., antibiotics, oxygen, ionic liquids, metals, organic pollutants, macromolecules, and nanomaterials) ^1–5^. Such defensive responses against environmental deterioration can improve the functional ability of cells and thus are crucial to the survival of biological species^3, 6–8^. Currently, increasing attention has been paid to the hormetic responses of bacteria exposed to antibiotics, given the close relationship to human and ecosystem health. For instance, antibiotics can dramatically alter the physiological functions of bacterial cells, even at concentrations far below lethal doses, which may contribute to human gut disorders and ecosystem dysfunction ^9, 10^.

At high concentrations, antibiotics are bacterial killers, whereas, at low concentrations, they can induce hormesis in various endpoints that may eventually favor the behavior of susceptible bacteria^11–13^. To date, a broad range of response endpoints for hormesis have been studied, including cell growth, secondary metabolic processes, and cellular functions, such as bioluminescence and biofilm formation ^14–17^. Some specific hormetic dose responses have been mechanistically explained, usually based on receptors and signaling pathways^16, 18^. For example, low doses of piperacillin can trigger secondary metabolite biogenesis via the *OxyR* and *SoxR* regulons in *Burkholderia thailandensis*^19^; neomycin and erythromycin can enhance bacterial bioluminescence in *Vibrio fischeri* via the *LuxR* quorum sensing (QS) system^20, 21^; and tetracycline can up-regulate type III secretion system expression and consequently biofilm formation in *Pseudomonas aeruginosa*^22^. Notably, although cell growth is one of the most common endpoints of hormesis, existing hormetic studies have mainly focused on microbial growth kinetics at hormetic concentrations of antibiotics^23–29^. However, the underlying molecular basis has yet to be clarified, probably because cell proliferation involves a complex array of responses, including both intrinsic and extrinsic cellular metabolic reactions.

Herein, we address this issue using *Comamonas testosteroni* bacteria, which are frequently present in diverse habitats, including activated sludge, marshes, marine habitats, plant and animal tissues, and the human gut^30–32^. Their diversified niches reflect effective adaptation to various physiochemical conditions and are thus a good model for exploring the molecular mechanism underlying hormesis. Sulfonamides (SAs) are the most extensively used antibiotics for bacterial treatment^33^. Importantly, no SA-related resistance genes (e.g., *sul1*, *sul2*, and *drfA*) were found in the complete genome of *C. testosteroni*^31^. We focused on the induction of cell proliferation by sulfamethoxazole (SMX) and sulfadiazine (SD) as a model for exploring how SAs stress induces hormetic effects. Follow the experimental procedure in Fig. S1, we provided a comprehensive understanding of the hormetic molecular mechanisms by which the LuxR solo transcription factor (TF) up-regulates fatty acid β-oxidation (FAO) through 3-ketoacyl-CoA thiolase (EC 2.3.1.16) under low-dose SAs stress (< 250 μg/L). Moreover, rapid cell metabolism accelerated the respiratory dissimilatory nitrate reduction to ammonium (DNRA) pathway in response to a reduced cellular redox state. Our results highlight the significant role of LuxR solo in regulating hormesis on cell growth as well as its generality in the Comamonadaceae family.

## Results

### Hormetic effect of SAs on *C. testosteroni* cell growth and morphology

The growth of *C. testosteroni* cells was examined upon the addition of 12 gradient concentrations (0 to 100 mg/L) of SAs (SMX, SD, and their mixture at 1:1 concentration). When cells reached the stationary growth phase (approximately 24 h), low-dose SAs (5 μg/L to 250 μg/L) significantly accelerated cell proliferation (reflected by OD_600_), whereas SAs at concentrations above 1 mg/L inhibited cell growth (Fig. 1a). Thus, these results demonstrated that SAs stress (SMX, SD, or their mixture) induced a hormetic effect on *C. testosteroni* cell proliferation. Notably, the hormetic effect elicited by the SAs mixture showed no significant difference from half the summation of the hormetic effects induced by separate SMX and SD exposure at all administered doses, except for 100 mg/L, suggesting that the functional groups in SMX and SD that induce these hormetic effects are identical (Table S1). Using flow cytometry to measure active biomass, we further confirmed the occurrence of the hormetic response of *C. testosteroni* under SAs stress, which started at approximately 12 h (1.09-fold, Wilcox-test *p* < 0.05) and peaked at 24 h (1.24-fold, Wilcox-test *p* < 0.05) (Fig. S2).

**Figure 1.**
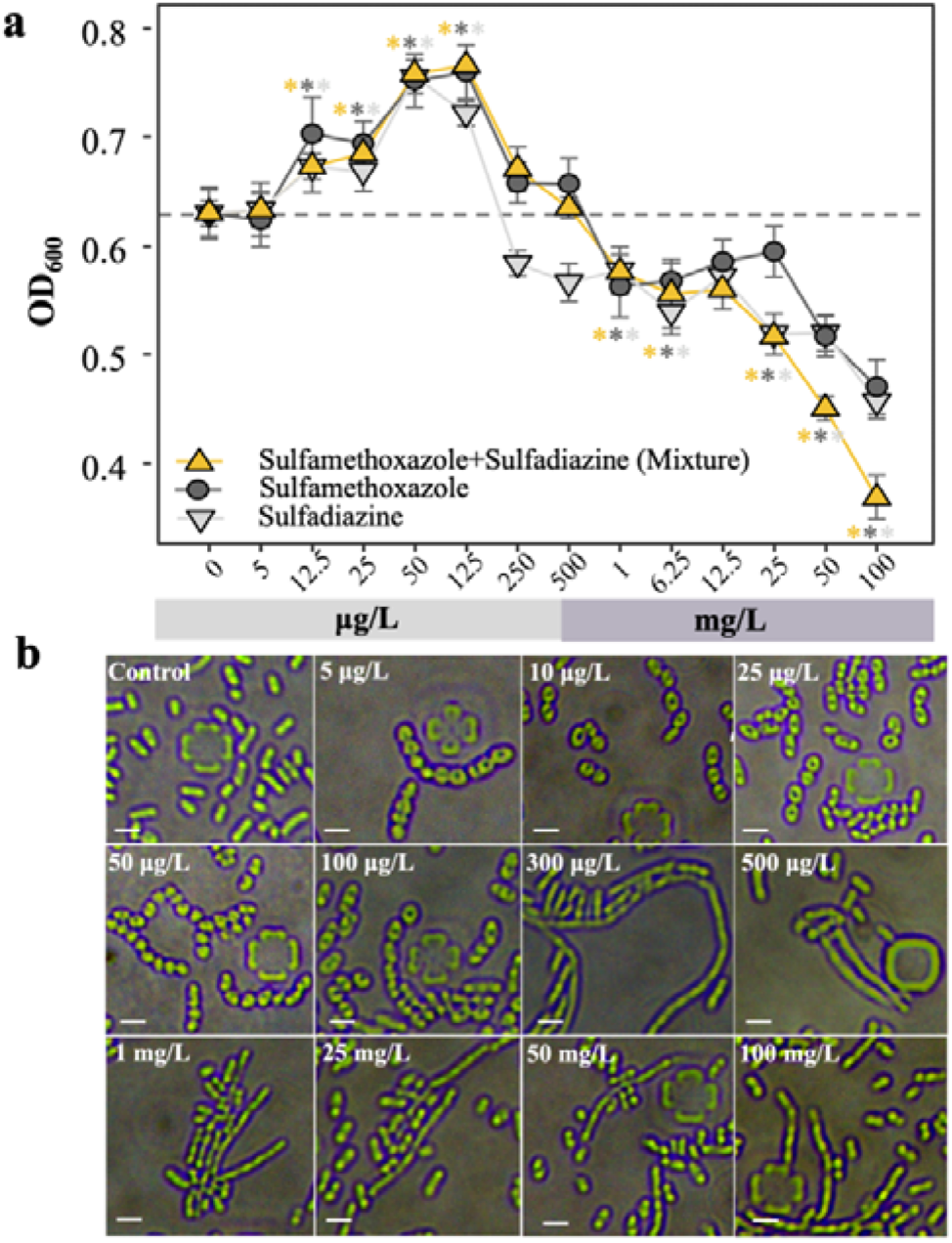
Hormetic effects of sulfonamides (SAs) on *C. testosteroni* cell growth and morphology. We first investigated the effects of ulfamethoxazole (SMX), sulfadiazine (SD), and their mixture (1:1 in concentration) on cell growth (**a**), represented as OD_600_ measured after 24 h of cell growth in a 96-well microplate containing MSM; mean and standard deviation of each result were calculated from six replicates. Dashed line represents average cell density without SAs treatment. Two-sample Wilcox test was used to evaluate significant differences in cell density between cultures with/without SAs treatment; \**p* < 0.05. We used inverted microscopy images to capture morphology variation of *C. testosteroni* after 24 h of cell growth inside the ONIX microfluidic platform containing MSM under different concentrations of SAs mixture (**b**). Brightfield images were acquired using a 100× oil immersion microscope objective. Scale bar, 2 µm.

Consistently, cell phenotypes changed along with the hormetic effect on cell growth. Under low-dose SAs stimulation (5 μg/L to 250 μg/L, including individual and mixed SAs), cells experienced a rod-to-ellipsoid transition and formed diplococci or short chains (Fig. 1b and Fig. S3). At the stationary phase (24 h) under SAs mixture exposure, mean cell length fluctuated slightly (0.54 ± 0.14 µm; n = 100) compared to the untreated cells (1.64 ± 0.15 µm) (Table S2). In contrast, under the inhibitory effect of high-dose SAs (> 500 µg/L), cells transitioned to very long filaments with normal cell widths (up to 10-fold the mean length of cells at the same growth stage without SAs treatment) (Fig. 1b and Fig. S3). Approximately 90% of cells were longer than 2.5 µm, more than 50% of which were > 5 µm (n = 100) (Table S2). Filamentary formation was due SA-induced DNA damage and replication perturbation, a phenomenon observed with SAs antibiotic treatment^34, 35^. However, the rod-to-ellipsoid shape alteration has not been reported previously. During growth, cells change their overall shape and size in rapid response to changing conditions by adjusting the synthesis of the cell envelope and cell division, which are tightly coordinated with DNA replication and protein synthesis through central metabolism^36^.

Together, these observations raise two questions: (i) what metabolic responses occur in *C. testosteroni* cells under the SA-triggered hormetic effect and (ii) how do SAs induce metabolic changes?

### SAs trigger metabolic adaptation responses in *C. testosteroni*

We next conducted transcriptomics analysis based on RNA sequencing (RNA-seq) of *C. testosteroni* under SAs mixture exposure to explore metabolic responses. Comparison between cells grown with/without 50 μg/L SAs mixture revealed a total of 486 significant differentially expressed genes (DEGs) (11.38% of all genes in the *C. testosteroni* genome), including 422 up-regulated DEGs. Comparison between cells grown with/without 1 mg/L SAs mixture revealed 673 significant DEGs (18.18% of all genes), including 378 up-regulated DEGs. Based on gene set enrichment analysis (GSEA) of differentially expressed proteins, cellular metabolism- and signal transduction-related biological processes were significantly up-regulated (Fig. 2a-c and Table S3).

**Figure 2.**
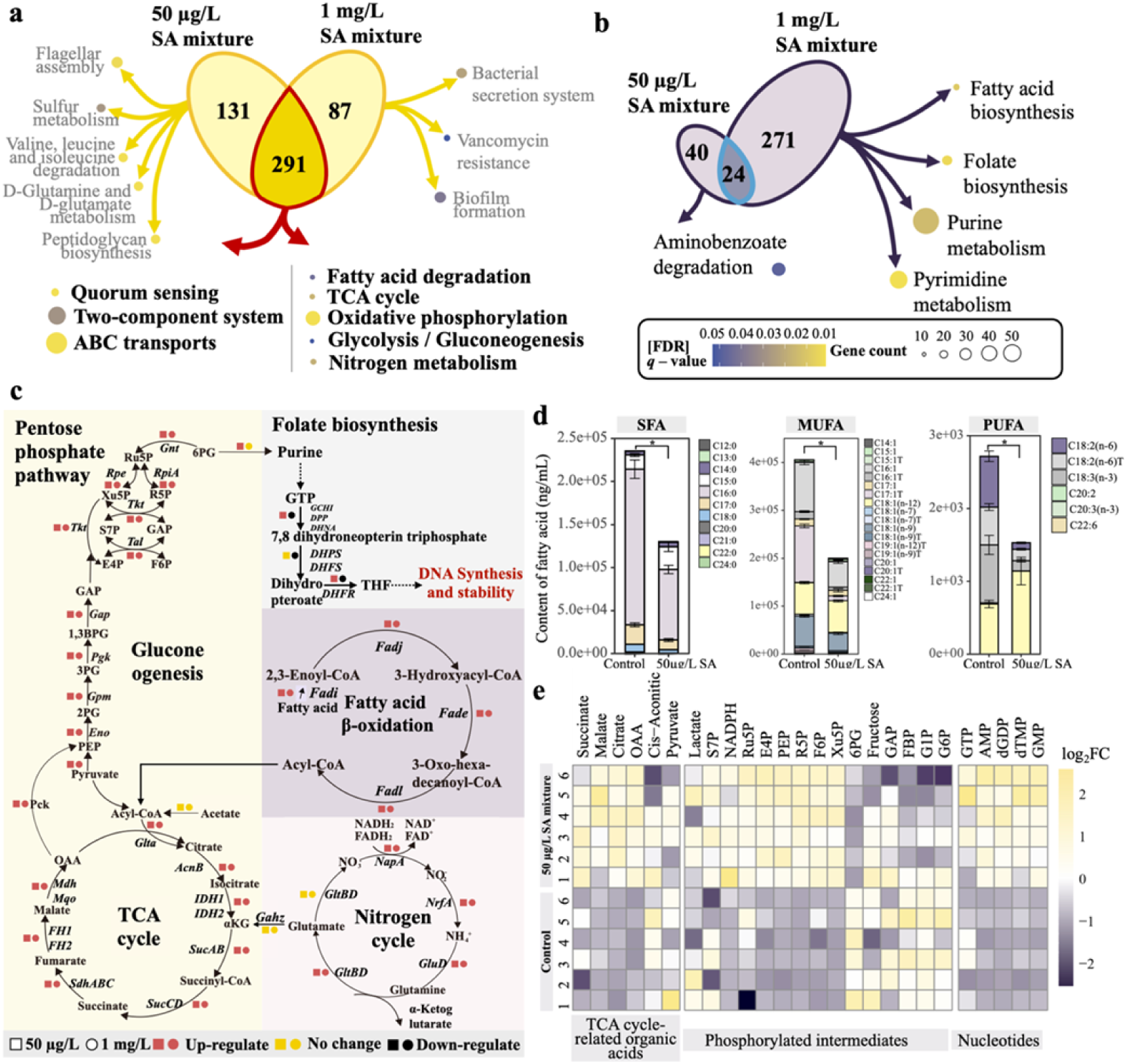
Global metabolic adaptation responses of *C. testosteroni* during growth under sulfonamides (SAs) mixture treatment. GSEA of differentially expressed proteins was performed to identify the up-regulated KEGG pathways (**a**) and down-regulated KEGG pathways (**b**) under SAs mixture treatment (50 μg/L and 1 mg/L, 1:1 concentration (mg/L) of sulfamethoxazole and sulfadiazine). In subfigure **(c)**, schematic shows metabolic regulatory network in *C. testosteroni* cells determined by transcriptomic analysis. Targeted LC-MS/MS-based metabolomics approach was further used to profile fatty acid (**d**) and central carbon (**e**) metabolites in *C. testosteroni* cells grown with/without 50 μg/L SAs mixture treatment. Significance (\**p* < 0.05) was determined using one-way analysis of variance (ANOVA) followed by Tukey HSD *post hoc* tests. **(d)** Intracellular pools (ng/mL) of SFAs, MUFAs, and PUFAs. Data are expressed as mean ± cumulative standard deviation of six biological replicates. (**e**) Heatmap shows fold-change (log_2_FC) in an intracellular pool of central carbon, divided into TCA cycle-related organic acids, phosphorylated intermediates, and nucleotides. Relative metabolite concentrations were normalized to a mean equal to 0 and standard deviation equal to 1. Metabolite abbreviations for Panels c and e are as follows: glucose-6-phosphate, G6P; fructose-6-phosphate, F6P; fructose-1,6-bisphosphate, FBP; dihydroxyacetone phosphate, DHAP; glyceraldehyde-3-phosphate, GAP; 6-phosphogluconate, 6PG; ribulose 5-phosphate, Ru5P; xylulose-5-phosphate, Xu5P; ribose-5-phosphate, R5P; sedoheptulose-7-phosphate, S7P; erythrose 4-phosphate, E4P; 1,3-biphosphoglycerate, 1,3BPG; 3-phosphoglycerate, 3PG; 2-phosphoglycerate, 2PG; phosphoenolpyruvate, PEP; oxaloacetate, OAA; α-ketoglutarate, αKG.

Among the catabolism categories, “fatty acid degradation” was the most significantly enriched biological process for both SA-tested concentrations (Fig. 2a and Table S3, n = 14, [false discovery rate (FDR)] *q*-value < 0.01), in which the activities of aldehyde dehydrogenase, acyl-CoA dehydrogenase, 3-hydroxy acyl-CoA dehydrogenase, and 3-ketoacyl-CoA thiolase were significantly up-regulated for both SA-tested concentrations (Fig. S4a, |log_2_FC| > 1.5, [FDR] *q*-value < 0.01). These enzymes are actively involved in FAO to generate NADH, flavin adenine dinucleotide (FADH_2_), acetyl-CoA, and NADPH ^37^. In the next few steps, the generated NADH and FADH_2_ enable the production of more adenosine triphosphate (ATP) (Fig. S5a), while acetyl-CoA enters the TCA cycle to produce citrate and other TCA cycle intermediates (Fig. S5b). Furthermore, NADPH can serve as a coenzyme for anabolic building blocks required for cell proliferation, such as lipid and nucleic acid synthesis^38^ (Fig. S5c). Increased carbon generated in the TCA cycle is further converted into phosphoenolpyruvate (PEP), which subsequently enters the gluconeogenesis and pentose phosphate (PP) pathways to provide additional substrates and energy for the biosynthesis of amino acids, lipids, and nucleotides. The enzymes involved in these processes were significantly up-regulated (Fig. S4b-d), especially the rate-limiting cataplerotic enzyme of the gluconeogenesis pathway, PEP carboxykinase (*Pck*) ^39^, which showed a 2.43-fold and 2.21-fold increase under 50 μg/L and 1 mg/L SAs mixture exposure, respectively (Fig. S4d). Compared with other nutrients, nucleotide production is of particular importance for proliferating cells, as it is needed to synthesize ribosomal RNA, duplicate the genome, and maintain the transcriptome^40^. Thus, we speculated that SAs may directly facilitate the oxidative decomposition of fatty acids to generate ATP, NADPH, and acetyl-CoA for further purine and pyrimidine biosynthesis, thus allowing *C. testosteroni* cells to synthesize sufficient nucleotides to support rapid cell growth. In contrast, cells exposed to the 1 mg/L SAs mixture showed a significant decrease in folate biosynthesis (Fig. 2b, Fig. S4f and Table S3, n = 10, [FDR] *q*-value < 0.01). This was expected because the bacteriostatic mechanisms of SAs can block the production of folic acid and therefore inhibit nucleotide synthesis^41, 42^(e.g., purine metabolism and pyrimidine metabolism), thus resulting in the growth inhibition effect observed at 1 mg/L SAs.

We next used a targeted metabolomics approach based on ultra-performance liquid chromatography-tandem mass spectrometry (UPLC-MS/MS) to validate the RNA-seq results. We profiled fatty acids and central carbon metabolites in *C. testosteroni* with/without 50 μg/L SAs mixture treatment. Targeted fatty acid quantification showed that SAs mixture exposure significantly decreased almost all oxidizable long-chain fatty acid content (Fig. 2d), including saturated fatty acids (SFA: C16-22:0), monosaturated fatty acids (MUFA: C15-24:1), and polyunsaturated fatty acids (PUFA: C18:2(n-6), C18:2(n-6) T, C18:3(n-3) and C20:3(n-3)). Furthermore, compared to growth without SAs treatment, the relative abundance of TCA cycle-related organic acids and phosphorylated intermediates was significantly increased under SAs treatment (Fig. 2e, 5.5-fold, *p* < 0.05). The cellular metabolite pools suggested that enhanced FAO decreased fatty acid content, thereby promoting carbon availability in the TCA cycle and accumulation of carbon in the upper gluconeogenesis pathway in SA-exposed cells, consistent with transcriptional pathway analysis.

Among the catabolism categories, the “nitrogen metabolism” biological process also exhibited significant enrichment (Fig. 2a and Table S3, n = 18, [FDR] *q*-value < 0.01). Most notably, the nitrate and nitrite reductases *Nap*/*Nrf*, which encode the respiratory dissimilatory nitrate reduction to ammonium (DNRA) pathway, were highly expressed under 50 μg/L and 1 mg/L SAs mixture exposure, respectively (Fig. S4e). It included the catalytic subunit *NapA* (2.24-fold and 2.19-fold), nonheme iron-sulfur cluster protein *NapG* (3.44-fold and 1.25-fold), pathway-specific chaperone *NapD* (2.48-fold and 1.12-fold), and pentaheme cytochrome c nitrite reductase *NrfA* (2.14-fold and 1.29-fold). The DNRA pathway produces energy (ATP) through oxidative phosphorylation via an electron transport chain (ETC)^43^, which can be used to maintain cellular activities and promote cell growth. Furthermore, this periplasmic pathway is also considered an effective electron sink to consume excess reduction forces, such as NADH and FADH_2_^44, 45^. Hence, the increased expression of respiratory DNRA genes in *C. testosteroni* cells growing with the SAs mixture may abrogate the oxidative stress induced by rapid cell proliferation. In addition, induction was not seen in genes associated with oxidative stress defence mechanisms (e.g., OxyR and SoxRS).

The quantitative reverse transcriptase polymerase chain reaction (qRT-PCR) results were consistent with the RNA-seq and metabolomic findings, confirming significant enrichment in fatty acid metabolism, biosynthesis, and DNRA pathways. Notably, select genes involved in FAO showed a 1.3–2.5-fold increase, PEP carboxykinase *Pck* showed a 1.4–2.0-fold increase, and *Nap/Nrf* systems showed a 1.3–2.2-fold increase (Fig. S6).

### SAs require transcriptional regulator LuxR solo to induce metabolic responses

The emerging view of metabolic regulation in proliferating cells is that signal transduction pathways and transcriptional networks participate in the significant reorganization of metabolic activities into a platform that supports bioenergetics, macromolecular synthesis, and, ultimately, cell division^46^. Therefore, we focused on the signal transduction-related category, in which “quorum sensing” was the most significantly enriched biological process for both SA-tested concentrations (Fig. 2a and Table S3, n = 21, [FDR] *q*-value < 0.01). Similar activation of QS has also been observed in studies on transcriptional modulation of bacterial gene expression using subinhibitory concentrations of antibiotics^47^. QS is a cell-cell communication system that uses exogenous signaling molecules and signal relay components to coordinate population density-dependent changes in behavior (e.g., morphogenetic and metabolic changes)^48^. We therefore speculated that low-dose SAs may act as a signaling molecule to activate QS, thereby regulating cell metabolic responses.

Among QS genes, the *LuxR* family transcriptional regulator (3.8-fold and 2.9-fold, [FDR] *q*-value < 0.01) and ABC transporter (1.8-fold and 1.6-fold, [FDR] *q*-value < 0.01) were highly expressed under 50 μg/L and 1 mg/L SAs mixture exposure, respectively (Fig. 3a). The qRT-PCR results further validated that the *LuxR* family transcriptional regulator was significantly induced (2.66–2.89-fold) under SAs treatment (Fig. S6). Sequence alignment and domain analysis of the identified LuxR transcriptional regulator (Table S4, accession no: WP_003076066.1) showed that the identified LuxR family protein was comprised of two complete functional domains, similar to typical QS proteins (Table S5), including a C-terminal region containing a predicted helix-turn-helix (HTH) motif implicated in DNA binding (e-value = 1.90e-11) and an N-terminal autoinducer-binding domain (e-value = 3.60e-12). Generally, an archetypical QS system in gram-negative bacteria is mediated by the LuxI and LuxR protein families^49^. LuxI-type proteins are N-acylhomoserine lactone (AHL) synthases that synthesize AHL signals, and LuxR family protein is regulator that directly bind to cognate AHL^49^. These protein-AHL complexes then bind to the specific gene promoter sequence to regulate the expression of the QS target gene^49^. Thus, we used BLASTP to identify proteins that may encode LuxI homologs. However, no unpaired or extra genes coding for LuxI homologs were present in the remainder of the genome. LuxR proteins with the same modular structure as QS LuxRs but devoid of a cognate LuxI AHL synthase are called solos^50^. Current evidence suggests that LuxR solo may exhibit a high degree of variability in both the type of ligands to which they respond and the mechanism by which they regulate target genes^51–53^. For example, the subfamily of LuxR solos in *Xanthomonas* no longer respond to endogenously produced AHLs but do respond to plant signals to participate in interkingdom communication between the host and pathogen^54^. Considering their potential ability to respond to various exogenous signals, we hypothesize that LuxR solo may provide the major pathway for SAs to induce metabolic responses. Thus, we next generated a *C. testosteroni* mutant based on *LuxR solo* gene knockdown to investigate the possible involvement of LuxR solo in the metabolic response (Fig. S7a and S7b). In the absence of SAs, the deletion of *LuxR solo* gene had no significant effect on cell growth (Fig. S7e). We then tested the ability of the deletion mutant Δ*LuxR* to trigger hormetic effects on cell growth under SAs mixture treatment. Notably, the Δ*LuxR* strains completely abrogated the SA-mediated growth induction effect and exhibited reduced cell density (up to 64%) despite low doses (< 250 μg/L). These effects were rescued by plasmid-encoded expression of *LuxR* in the ΔLuxR strain (Fig. 3b, Fig. S7c-e). These results indicate that LuxR solo is required for SA-mediated induction of metabolic responses. Based on these findings, we proposed a mechanistic hypothesis of hormesis:

- At low doses, SAs are first completely bound by membrane protein LuxR solo, allowing LuxR solo-SA complexes to accumulate. After 12 h, the accumulated LuxR solo-SA complexes bind to the target gene and activate (i) FAO to provide additional nutrients for biosynthesis and/or (ii) DNRA to modulate ATP formation and redox balance (Fig. 3c).
- As the dose increases, the binding reactions reach equilibrium when the stimulating effect reaches saturation. However, free-SAs begin to bind to dihydrofolate reductase, thereby inhibiting the synthesis of folate and causing an inhibitory effect on nucleic acid synthesis. The inhibitory effect increases with SAs dose, and when inhibition surpasses stimulation, the total effect of inhibition is observed.

**Figure 3.**
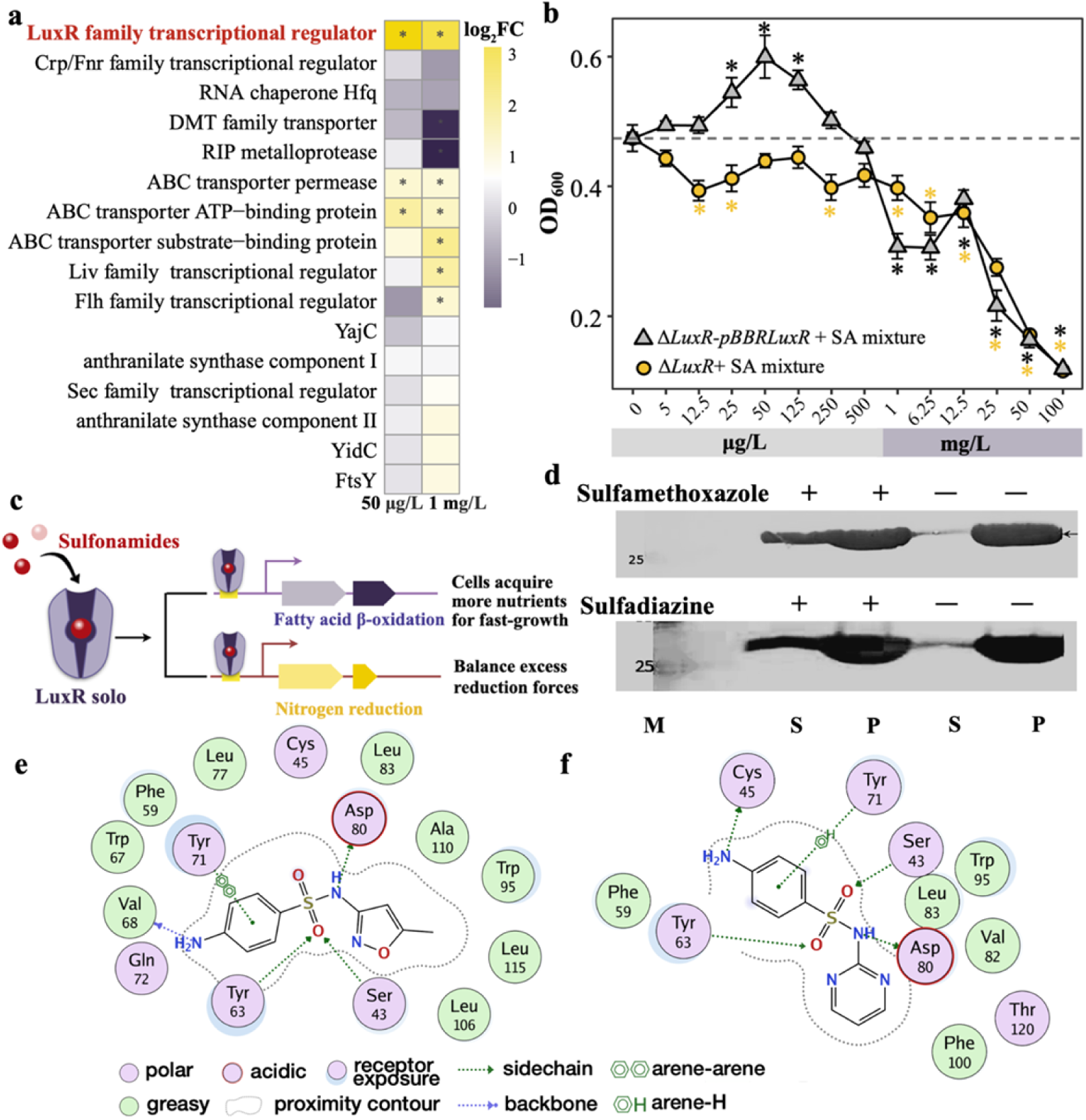
Sulfonamides (SAs) require transcriptional regulator LuxR solo to induce metabolic responses. **(a)** Heatmap shows fold-change in gene expression (log_2_FC) in QS pathway under SAs mixture treatment (50 μg/L and 1 mg/L, 1:1 concentration (mg/L) of sulfamethoxazole (SMX) and sulfadiazine (SD)); \**p* < 0.05. Red represents screened potential SA signal transduction gene LuxR solo (|log_2_FC| >1.5, [FDR] *q*-value < 0.01). **(b)** LuxR solo is required for SA-mediated metabolic responses, as determined using *C. testosteroni* mutants Δ*LuxR* and Δ*LuxR-*pBBR*LuxR* to trigger hormesis effect on cell growth [OD_600_] under SAs. Dashed line represents average cell density without SAs treatment. Wilcox test was used to evaluate significant differences in cell density [OD_600_] between cultures with/without SAs treatment. Significance corresponds to adjusted Wilcox-test *p*-values (\**p* < 0.05). (**c**) Combining subfigures (a) and (b), we show a proposed schematic overview of metabolism regulation by the LuxR family transcriptional regulator. Subfigures **(d)-(f)** verify the direct interaction of LuxR solo protein and SAs based on solubility measurements of overexpressed LuxR solo protein and docking studies. **(d)** Western blotting was used to show LuxR solo in soluble (S) and pellet fractions (P) of cell lysates from recombinant *E. coli* strain BL21(DE3) supplied with 75 µM SMX or SD. M denotes marker (representative bands are labeled). Docking view of SMX **(e)** and SD **(f)** at LuxR solo binding site. Green dashed lines indicate hydrogen bonds between ligand and amino acid residues.

### SAs directly bind to LuxR solo via sulfonyl groups to regulate target pathways

To test the first mechanistic hypothesis, we investigated the binding of SA molecules to LuxR solo proteins. Generally, QS-LuxR proteins are unfolded, proteolyzed, or form inclusion bodies in the absence of a cognate molecule, whereas in the presence of and when bound to cognate molecules, they become folded and soluble^55–57^. Therefore, we used this biochemical “folding switch” feature to explore whether LuxR solo can directly interact with SAs. We reasoned that if SAs act as AHL-like molecules, the expression of the protein in the presence of SAs will primarily produce soluble proteins. We cultured *E. coli* BL21(DE3) overexpressing His-LuxR in the absence and presence of SMX and SD and identified the presence of soluble LuxR solo using western blot analysis. When LuxR solo was overexpressed in the absence of SAs, almost all the LuxR solo protein was found in the particulate fraction, indicating that it accumulated as insoluble inclusion bodies. In contrast, the addition of 75 mM SMX or SD increased LuxR solo solubility, with approximately half of the total LuxR solo protein found in the soluble fraction (Fig. 3d). These results indicate that LuxR solo proteins may be involved in combination with SAs.

Molecular docking was further performed to visualize the binding process. First, the SWISS-MODEL server was used to successfully generate a 3D structure for the LuxR solo protease using the crystal structure of SdiA in complex with 3-oxo-C6-homoserine lactone (PDB ID: 4Y15) as the template. Both SMX and SD were then docked onto the template. Results showed that SMX and SD were stable in the binding pocket of LuxR solo protease. The minimum binding free energy of SMX with the LuxR solo protein was −7.89 kcal mol^−1^, stabilized in the binding pocket by four hydrogen bonds, i.e., between the sulfonyl group and residues Ser-43 and Tyr-63 and between the imino/amino group and residues Asp-80 and Val-68. In addition, a π-π stacking interaction existed between the benzene ring of SMX and the alkyl atoms of Tyr-71 (Fig. 3e and Fig. S8). Similarly, the minimum binding free energy with SD was −7.40 kcal mol^−1^, with Asp-80, Cys-45, Tyr-63, and Ser-43 serving as anchoring points for the ligands (Fig. 3f and Fig. S9). Tyr-63 and Ser-43 formed hydrogen bonds with sulfonyl groups, Asp-80 and Cys-45 formed hydrogen bonds with the imino/amino groups, and Tyr-71 formed a C-H bond with the benzene ring. As sulfonyl groups were crucial in binding LuxR solo with SMX or SD, we docked 14 other sulfonyl-containing chemicals with the LuxR solo template using the same docking process. Among the compounds that induced hormetic effects after 24 h of growth, all showed binding free energies below −5.14 kcal mol^−1^ (Fig. S10). These results suggest that *C. testosteroni* hormesis is strongly influenced by the interaction between LuxR solo and sulfonyl functional groups.

### LuxR solo regulates 3-ketoacyl-CoA thiolase to activate fatty acid β-oxidation

Due to the presence of the LuxR-type HTH motif in the C-terminal domain and in agreement with the HTH position/function relationship postulated by Perez-Rueda et al.^58^, we postulated that the LuxR solo-SA complex may bind to the target gene promoter region. Therefore, we next identified the direct functional target of LuxR solo-SA in *C. testosteroni*. We performed RNA-seq of the *C. testosteroni* mutant Δ*LuxR* treated with 50 µg/L SAs mixture and determined that LuxR solo regulated the fatty acid oxidation and/or nitrogen metabolism pathways, although other activation pathways could not be ruled out. We defined the target gene of LuxR solo based on the following criteria: (i) the gene significantly expressed in wild-type (WT) *C. testosteroni* cells was no longer changed after deleting *LuxR solo* gene upon low-dose SAs treatment and (ii) its upstream promoter region contained the *lux-box* bound by LuxR solo. Nine clusters associated with FAO and nine clusters associated with DNRA were investigated. We observed that DNRA genes in the deletion mutant Δ*LuxR* were up-regulated, a transcriptional pattern similar to that observed in SA-exposed WT cells (Fig. S11). In contrast, the FAO gene clusters that showed significant changes in WT cells disappeared in the Δ*LuxR* mutants (Fig. 4a). Analysis based on qRT-PCR further validated the RNA-seq results, showing that only significant changes in the fatty acid oxidation pathway were lost in mutant cells (Fig. S12). Thus, these results suggest that LuxR solo may potentially regulate the FAO pathway but not the DNRA pathway.

**Figure 4.**
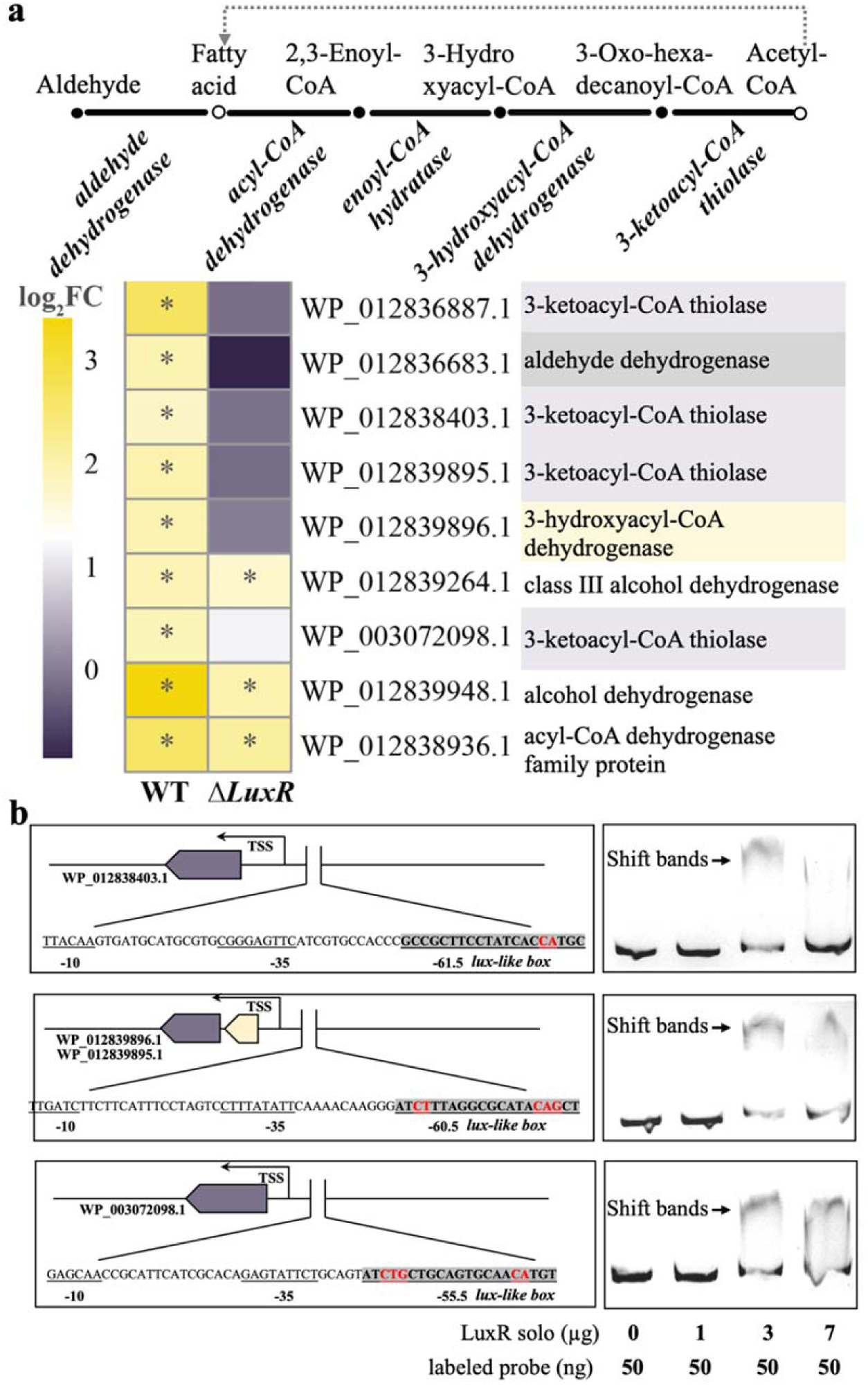
LuxR solo directly up-regulates fatty acid β-oxidation. Subfigure **(a)** summarizes fatty acid β-oxidation pathway (top), and heatmap (below) shows related fold-change in gene expression (log_2_FC) in wild-type and mutant Δ*LuxR* upon low-dose sulfonamides (SAs) mixture treatment (50 μg/L, 1:1 concentration). *P*-values are compared to strains without SAs treatment (\**p* < 0.05). Three binding sites (*lux-like box*) were identified in the 3-ketoacyl-CoA thiolase promoter region **(b)**. Sequences of putative LuxR binding sites (left) are shown in the 5′ to 3′ direction (gray); red represents conserved binding sites. EMSA (right) shows that LuxR solo binds to 3-ketoacyl-CoA thiolase promoter region. Conditions are indicated below each lane.

To further identify the direct binding site of LuxR solo, we focused on three operons involved in FAO, aldehyde dehydrogenase (WP_012836683.1), 3-hydroxyacyl-CoA dehydrogenase (WP_012839896.1), and 3-ketoacyl-CoA thiolase (WP_012836887.1, WP_012838403.1, WP_012839895.1, and WP_003072098.1). *LuxR* family regulator is reported to control target genes by binding to a conserved 20-bp *lux-box* CTG-(N10)-CAG sequence in the gene promoter, in which five of the six bases are essential for LuxR binding^59^. Therefore, we retrieved the conserved *lux-box* in the promoter region of six FAO genes using MEME tools^60^. Results showed that putative *lux-like box* motifs were only identified in the promoter regions of 3-ketoacyl-CoA thiolase (WP_012838403.1, WP_012839895.1, and WP_003072098.1), located −61.5, −60.5, and −55.5 bp upstream of the transcription start sites (TSS) (Fig. 4b, left), respectively. As noted previously, *LuxR* in *V. fischeri* is a transcriptional activator rather than a repressor when the position of the *lux-box* is located −60 bp from the center to the TSS. This suggests that the LuxR solo protein identified here is also a transcriptional activator that interacts with the C-terminal domain of the alpha subunit of RNA polymerase (NAP; subunit composition α2ββ′σ)^61^.

To test whether LuxR solo can directly bind to the 3-ketoacyl-CoA thiolase promoter region, we performed an electrophoretic mobility shift assay (EMSA). Using the 670-bp, 570-bp, and 600-bp promoter sequences of 3-ketoacyl-CoA thiolase (WP_012838403.1, WP_012839895.1, and WP_003072098.1), we observed a LuxR solo-dose-dependent mobility shift, consistent with the formation of a DNA-protein complex (Fig. 4b, right). This suggests that the QS regulator LuxR solo can directly activate the expression of 3-ketoacyl-CoA thiolase.

### Proposed regulatory pathway for hormetic effect

Based on the above experiments, we propose that the entire regulatory pathway (Fig. 5) is as follows: Low-dose SAs (< 250 μg/L) interact with the LuxR solo autoinducer-binding site via the sulfonyl group, forming a LuxR solo-SA complex. The complex then triggers the transcription of degradative thiolase [3-ketoacyl-CoA thiolase (EC2.3.1.16)], which catalyzes the last reaction of the FAO cycle to regenerate acyl-CoA for another round of β-oxidation and release of acetyl-CoA for the citric acid cycle^62^, thus activating cell proliferation. Along with this metabolic boost, unavoidable byproducts of aerobic metabolism, such as reactive oxygen species from NADH, may damage cell redox homeostasis. Our results showed that the cytoplasmic pathway for NO_3_^−^ and NO_2_^−^ reduction involving periplasmic reductases *Nap* and *Nrf* was simultaneously enhanced to maintain redox balance in proliferating cells.

**Figure 5.**
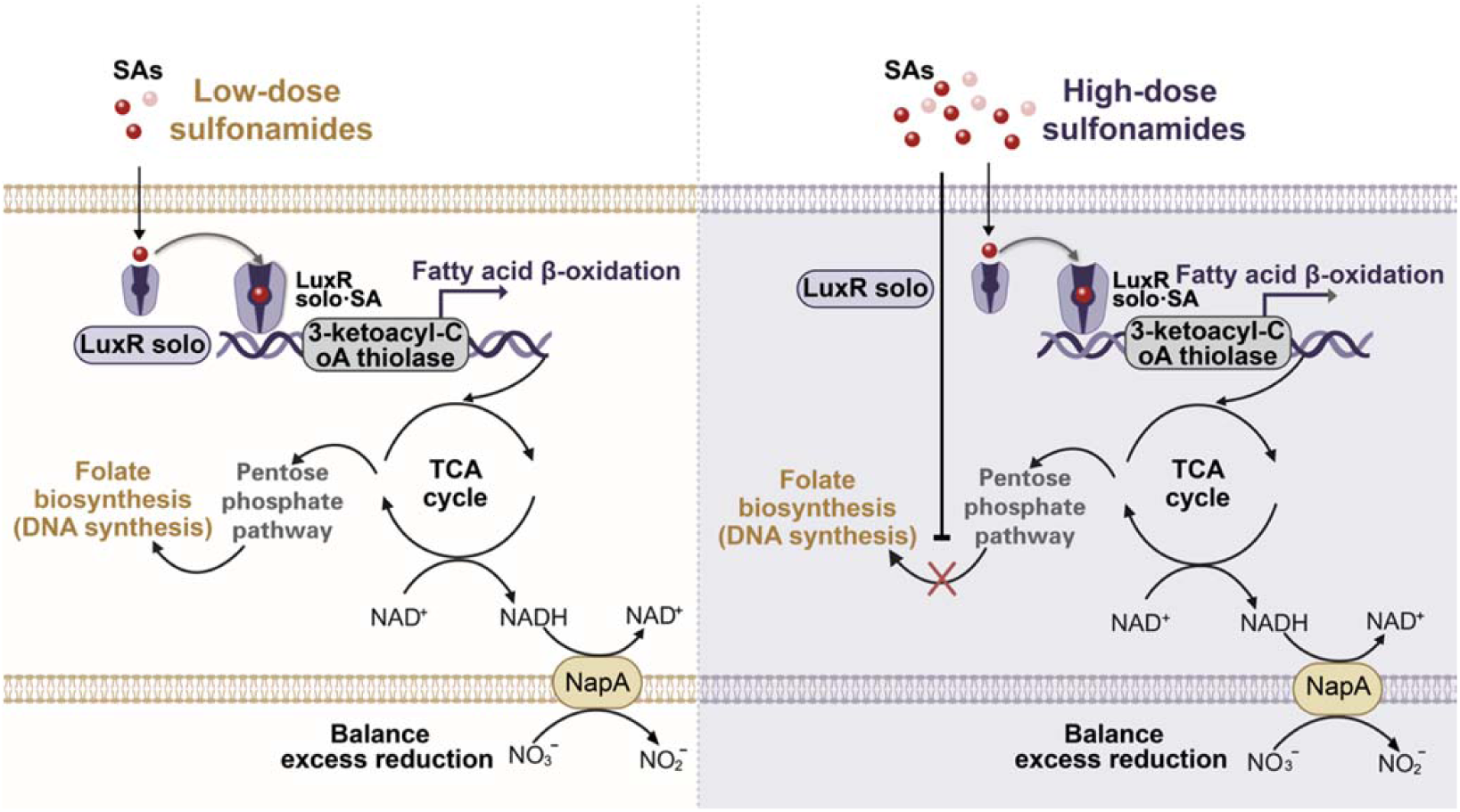
Proposed regulatory pathway of *C. testosteroni* hormesis under sulfonamides (SAs). At low exposure, SAs can bind to LuxR solo and cause accumulation of LuxR solo-SA complexes. Accumulated LuxR solo-SA complexes then activate fatty acid β-oxidation in low dose-exposed bacteria, thus exhibiting hormesis of cell growth. As doses increase at high exposure, more SAs bind to dihydropteroate synthase and inhibit folate biosynthesis. When inhibition surpasses stimulation, toxicity is observed.

### Generality of regulatory pathway for hormetic effect

We blasted the TF LuxR solo (WP_003076066.1) in the NCBI database and found that LuxR solo protein was conserved among Comamonadaceae family members, including the genera *Comamonas*, *Acidovorax*, and *Variovorax*^63^ (Fig. 6a and Fig. S13). We randomly selected four strains from the latter two genera, i.e., *Acidovorax avenae*, *Acidovorax delafieldii*, *Variovorax paradoxus*, and *Variovorax soli*, for culture under SAs stress. All stains established significant SA-induced hormetic effects on cell growth (Fig. 6b), supporting the generality of the regulatory model in the Comamonadaceae family.

**Figure 6.**
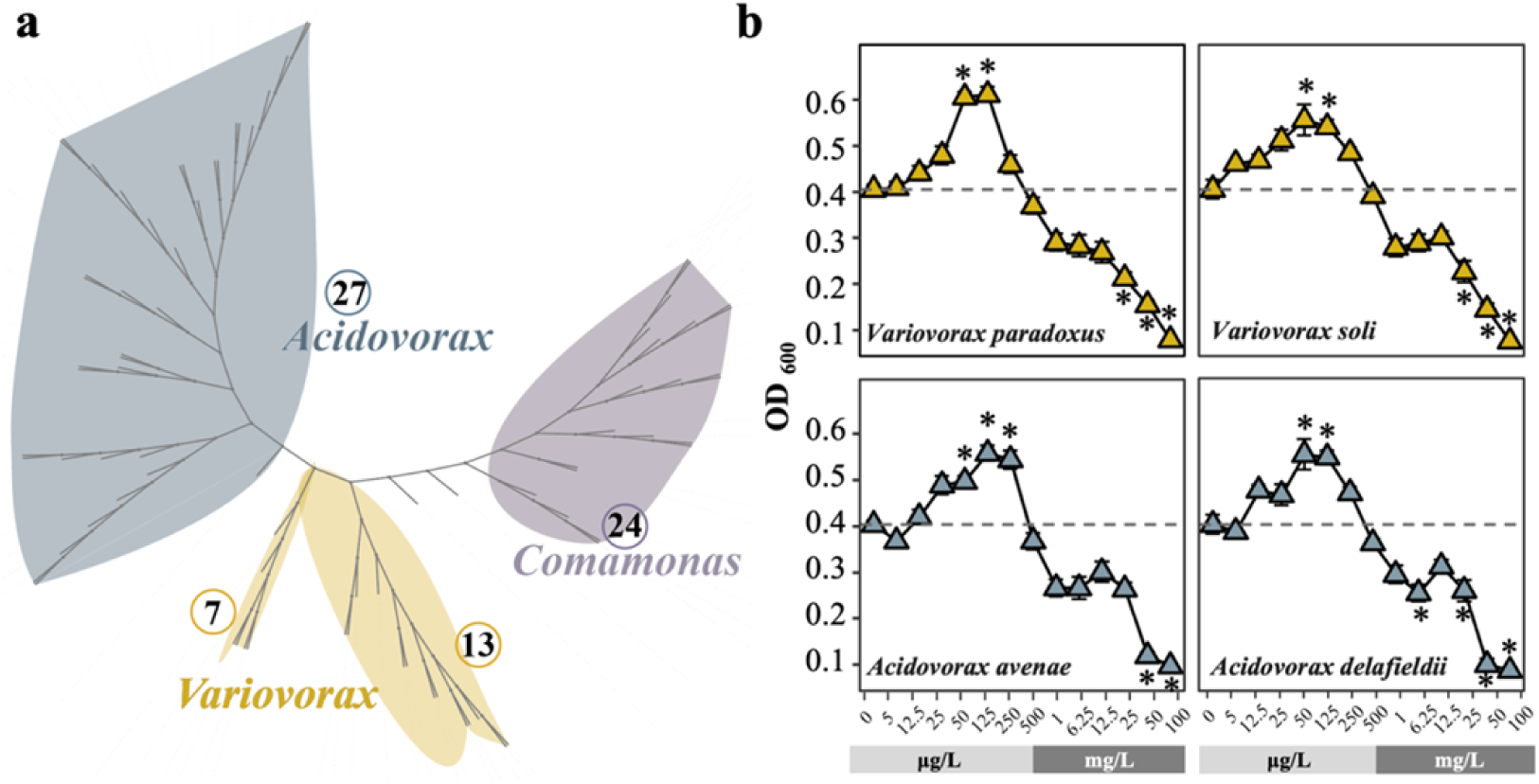
LuxR solo is conserved in the Comamonadaceae family, including the genera *Comamonas*, *Acidovorax*, and *Variovorax* (**a**). Numbers represent species counts in each genus that contain the LuxR solo protein. (**b**) Hormetic effects on cell growth in *Variovorax paradoxus*, *Variovorax soli*, *Acidovorax avenae*, and *Acidovorax delafieldii.* OD_600_ was measured after 24 h of cell growth in a 96-well microplate containing MSM, stains, and different concentrations of sulfamethoxazole and sulfadiazine mixtures (1:1 in concentration); mean and standard deviation of each result were calculated from 12 replicates. Dashed line represents average cell density without sulfonamides (SAs) treatment. Wilcox test was used to evaluate significant differences in cell density between cultures with/without SAs treatment. Significance (\**p* < 0.05) was determined using Wilcox test.

## Discussion

From an eco-evolutionary perspective, the ability to adapt to antibiotics is a trait that has evolved in single-celled organisms living in natural environments^64^. While antibiotics have traditionally been perceived as weapons that can inhibit bacterial growth, recent work also emphasizes their role as concentration-dependent signaling molecules^65–67^. In the current study, we showed that SAs function as signaling molecules that can modulate cell physiology and phenotype of bacteria at lower concentrations.

In response to SAs stress, *C. testosteroni* cells undergo adaptive metabolic reprogramming characterized by the broad activation of FAO (Fig. 2c). Cell metabolism can be perceived as a complex network of pathways with plasticity, feedback loops, and crosstalk to ensure cell fitness^68^. Plasticity is crucial and may be partially provided by FAO, which generates additional ATP and NADPH, eliminates potentially toxic lipids, and provides metabolic intermediates for cell growth. The ability of cells to grow and survive is largely limited by the levels of cytosolic NADPH, a coenzyme of anabolic enzymes^69^. Therefore, metabolic reprogramming must meet the requirement of producing reduced NADP^+^^70^. Generally, this is achieved through three primary enzymatic reactions, i.e., oxidation of glucose via the PP pathway, metabolism of malate to pyruvate, and oxidation of isocitrate to α-ketoglutarate. However, *C. testosteroni* strains lack the genes associated with G6P dehydrogenase in the oxidative PP pathway ^71, 72^. Evolutionarily, this branch of the PP pathway is a newer metabolic strategy due to its absence in many thermophilic organisms, archaea, and aerobic bacteria^73^. The lack of the oxidative branch of the PP pathway leads to the production of NADPH only from the TCA cycle, which is insufficient to support anabolism in rapidly proliferating cells^74^. Previous studies have proposed that *C. testosteroni* may leverage transhydrogenase conversion of NADH to NADPH to meet the required NADPH flux^74^. Our results demonstrated that under SAs stress, increased FAO-dependent NADPH may compensate for the deficit in NADPH production in the oxidative PP pathway and thus meet the biosynthetic requirements of proliferating cells under low-dose SAs.

Our study showed that the enhanced FAO process decreased the content of intracellular fatty acids (Fig. 2d). Evidence suggests that fatty acids are a positive determinant of cell size due to the limited fatty acid pool available for phospholipid synthesis^75, 76^. For example, curtailing fatty acids reduces the size of gram-negative *E. coli*, gram-positive *Bacillus subtilis*, and unicellular *S. cerevisiae* ^69, 75, 77^. Our findings support the concept that cell size is dictated by the capacity of the cell envelope, which is a product of nutrient-dependent changes in fatty acid availability. Cell size must be coordinated with other biosynthetic processes to maintain cell envelope integrity to link surface area expansion with increases in cytoplasmic volume. Our results showed that with decreasing fatty acid availability under low-dose SAs stress, cells morphed into an ovoid shape at the lowest surface-to-volume ratio.

In summary, under low-dose SAs stress, intrinsic and extrinsic molecular mechanisms lead to alterations in core cellular metabolism processes to support the three basic needs of proliferating cells, i.e., rapid ATP generation to maintain energy status, increased biosynthesis of macromolecules, and tightened maintenance of appropriate cellular redox status. Conversely, high-dose SAs stress has antimetabolite effects that induce cell death via inhibition of dihydrofolate reductase^35^. The hormetic response, therefore, is governed by partial versus complete inhibition of the target and associated consequences. Partial inhibition allows bacteria to reorganize metabolic pathways (i.e., enhanced FAO pathway) to mount a counterattack. In contrast, complete or near-complete inhibition drains resources, leading to cell death. Overall, our findings provide novel mechanistic insight into the genes and enzymes involved in the hormetic effect of antibiotics in bacteria. Importantly, the inherent ability of *Comamonas* species to survive in ecological niches and the human gut makes them formidable candidates to cause mild but persistent infections ^78^, especially in individuals with predisposing conditions.

### Limitations of the study

Although we identified LuxR solo induction of FAO as a major mechanism underlying *Comamonas* hormesis, other pathways may also exist given that *LuxR* homologs can crosstalk with other cytoprotective TFs to mediate group behavior^79^. Therefore, further work is needed to explore the potential link between LuxR solo and other TFs in *Comamonas* hormesis.

## Materials and Methods

### Bacterial strains and culture

Bacterial strain growth experiments were conducted using microplate reader kinetic assays. For *C. testosteroni* CNB-2, overnight cultures were inoculated in 50 mL of nutrient broth (NB) medium at 30°C/200 rpm. These cultures (OD_600_ ≈ 0.1) were then washed twice with phosphate-buffered saline (PBS) buffer and inoculated at a 3% inoculation proportion (v:v) into 200 μL of mineral salt medium (MSM) (Table S6) containing potential hormesis inducers in 96-well plates. The potential inducers included 14 sulfonyl-containing compounds (structures and properties are listed in Table S7). The plates were sealed, and optical density (OD) was monitored using a Spark™ 10 M microplate reader (Tecan, Switzerland) at 600 nm (OD_600_) with fast continuous shaking. *Escherichia coli* strains DH5αλpir, β2155, and WM3064 were used to construct the *C. testosteroni* CNB-2 mutants, and strain BL21(DE3) was used for protein expression. The *E. coli* cells were grown in sterile NB broth at 37°C/200 rpm using 25 μg/mL gentamycin and 50 μg/mL ampicillin for the corresponding mutants. All bacterial strains, gene deletion mutants, and plasmids used in this study are listed in Table S8.

### Microfluidics

The CellASIC ONIX Microfluidic Platform (Merck Millipore, Germany) was used to maintain cells growing in a monolayer to monitor growth behavior under SAs stress. Cell inoculum (50 μL; OD_600_ ≈ 0.1) was loaded onto a microfluidic plate B04A-03 at 4 psi for 15 s. PBS was then added at 1 psi for 30 s to remove non-trapped cells. Subsequently, 250 µL of MSM containing different concentrations of SAs was pipetted into the wells and incubated under a total pressure of 2 psi (flow rate of 10 µL/h) for 72 h. In all cases, images were taken using a MshOt microscope equipped with an MD3 digital camera (MshOt Microscopy Imaging Expert; Guangzhou, China) and analyzed using an MshOt Digital Microscope Imaging System v1.0. Time-lapse images were captured at 1-h intervals for 48 h. A bright-field image was acquired at each time point to evaluate growth behavior.

### Construction of *C. testosteroni* CNB-2 mutants lacking *LuxR*

All primer pairs used for gene knockout are listed in Table S9. Upstream and downstream genes flanking *LuxR* were amplified by PCR using two primer sets, i.e., LuxR-5F/LuxR-5R and LuxR-3F/LuxR-3R. The gentamicin resistance gene (*Gm*) gene, which confers resistance to gentamycin, was amplified from the plasmid pJQ200SK by PCR using the primer set Gm-F and Gm-R. The upstream fragment, *Gm,* and downstream fragment junction was amplified via fusion PCR and cloned into pCVD442, a suicide plasmid containing the ampicillin resistance gene, to obtain recombinant plasmids. Then, pCVD442 containing the *LuxR* gene was introduced into the *E. coli* strain β2155 via electroporation to obtain donor plasmids and introduced into the recipient CNB-2 strain via conjugation. Transconjugants were selected on NB plates supplemented with ampicillin (50 μg/mL) and gentamycin (25 μg/mL). The *C. testosteroni* CNB-2/Δ*luxR* mutants were confirmed by PCR and sequencing using two primer sets, i.e., LuxR-outF/LuxR-outR and LuxR-inF/LuxR-inR.

### Complementation mutants carrying *LuxR*

All primer pairs used for gene complementation are listed in Table S9. To construct the mutant complemented with *C. testosteroni LuxR*, we amplified the *C. testosteroni LuxR* gene by PCR using LuxR-comF and LuxR-comR primers. The In-Fusion technique was used to clone the DNA fragment into the EcoRV site of the kanamycin-resistant plasmid vector pBBR1MCS2 to yield pBBR1MCS2-*LuxR*. The plasmid was then transformed into the *E. coli* strain WM3064 via electroporation and conjugation with the Δ*luxR* mutant. Transconjugants were selected on NB plates containing kanamycin (50 µg/mL). The *C. testosteroni* CNB-2/Δ*luxR-*pBBR*LuxR* mutant was confirmed by PCR using the pBBR1-F and pBBR1-R primers.

### RNA-seq analysis

RNA-seq was used to quantify transcriptional abundance of the *C. testosteroni* CNB-2 wild-type, Δ*LuxR*, and Δ*luxR-*pBBR*LuxR* strains under different SAs stresses: (i) WT cells was cultured for 24 h at 30°C/200 rpm with/without SAs treatment (50 μg/L and 1 mg/L, mixture of SMX and SD), three groups; (ii) Δ*LuxR* cells were cultured for 24 h at 30°C/200 rpm with/without SAs treatment (50 μg/L, mixture of SMX and SD); and (iii) Δ*LuxR-*pBBR*LuxR* cells cultured for 24 h at 30°C/200 rpm without SAs treatment (50 μg/L, mixture of SMX and SD). All strains were cultivated in suspension cultures overnight and conducted in triplicate. Total RNA was extracted using a Qiagen mini-RNA prep kit (Qiagen, Germany) following the manufacturer’s instructions. Extracted RNA was kept at −80°C before cDNA library construction. Total RNA concentration, RNA integrity number (RIN), and RNA quality number (RQN) were evaluated using an Agilent 2100 Bioanalyzer (Santa Clara, USA). Samples with an RIN/RQN value above 8.0 were collected for sequencing. Paired-end sequencing was performed on the Illumina HiSeq 2500 platform with a read length of 150 nucleotides (San Diego, CA, USA). The raw RNA-seq data were deposited in the NCBI GEO Short Read Archive (SRA) under accession numbers PRJNA933284. Sequencing reads were assembled and analyzed using the NCBI Prokaryotic Genome Annotation Pipeline with the reference CNB-2 strain genome (GenBank: CP001220.2). Data were normalized by calculating fragments per kilobase per million mapped fragments (FPKM). Significant changes in gene expression were defined based on ≥1.5-fold-change in FPKM and [FDR] *q*-value < 0.01.

### Quantitative real-time polymerase chain reaction (qRT-PCR)

Under treatment with SMX (50 μg/L), SD (50 μg/L), and their mixture (50 μg/L, 1:1 concentration), the differential expression levels of nitrogen metabolism genes (*NapA*, *NapD*, *NirD*, *NrfA*, and *GluD*), fatty acid β-oxidation genes (*Fadi*, *Fadj*, *Fade*, and *Fadl*), biosynthesis gene *Pck*, and QS gene *LuxR solo* were validated using qRT-PCR. To perform reverse transcription analysis, RNA samples were used as templates to synthesize cDNA using a cDNA synthesis kit (Qiagen, Germany). The resultant cDNA was then used for qRT-PCR using the LightCycler 96 Real-Time PCR system (Roche Diagnostics, Switzerland). The relative expression level (copy number of mRNA transcript) of each target gene was normalized to the cDNA concentration and compared with control samples (no SAs treatment). All primer pairs used for qRT-PCR analysis are listed in Table S9.

### Targeted metabolomics analysis with LC-MS/MS

#### (i) Fatty acids

The *C. testosteroni* WT and Δ*luxR* mutant cells (∼10^8^) cultured with/without SAs mixture were homogenized with 300 μL of isopropanol/acetonitrile (1:1) containing mixed internal standards and centrifuged at 4°C/12 000 rpm for 10 min. The supernatant was then injected into the LC-MS/MS system for analysis. Ultra-high-performance liquid chromatography-tandem mass spectrometry (UHPLC-MS/MS) (ExionLC™ AD UHPLC-QTRAP 6500+, AB SCIEX Corp., Boston, MA, USA) was used to quantify fatty acids at Novogene Co., Ltd. (Beijing, China). Separation was performed on a Waters Acquity UPLC BEH C18 column (2.1 × 100 mm, 1.7 μm) maintained at 50°C. The mobile phase, consisting of 0.05% formic acid in water and isopropanol/acetonitrile (1:1), was delivered at a flow rate of 0.30 mL/min. The ratio of the concentration of the standard to the internal standard was used as the abscissa, and the ratio of the peak area of the bar to the internal standard was used as the ordinate to investigate standard solution linearity. The limit of quantification (LOQ) was determined by the signal-to-noise ratio (S/N), which compares the signal measured by the standard solution concentration with the blank matrix. All fatty acid substances were tested, and their categories are listed in Table S10.

#### (ii) Central carbon

The *C. testosteroni* WT and Δ*luxR* mutant cells (∼10^8^) cultured with/without SAs mixture were individually ground with liquid nitrogen. The homogenate was resuspended in 500 μL of prechilled 80% methanol and 0.1% formic acid in a vortexing well. The samples were incubated on ice for 5 min, then centrifuged at 15 000 rpm and 4°C for 10 min. Aliquots of the supernatant were diluted to a final solution containing 53% methanol using LC-MS grade water. The samples were transferred to a fresh Eppendorf tube and centrifuged at 15 000 × *g* and 4°C for 20 min. Finally, the filtrate was injected into the UHPLC-MS/MS system for analysis. Each experimental sample was taken in equal volume and blended as quality control (QC) samples. The blank sample was a 60% methanol aqueous solution containing 0.1% formic acid instead of the experimental sample, and the pretreatment process was the same as the experimental sample.

For UHPLC-MS/MS analysis, a QTRAP® 6500+ mass spectrometer was operated in positive polarity mode with a curtain gas of 35 psi, collision gas of medium, ion spray voltage of 4 500 V, temperature of 550°C, ion source gas of 1:60, and ion source gas of 2:60. A negative ion mode QTRAP® 6500+ mass spectrometer was operated in negative polarity mode with a curtain gas of 35 psi, collision gas of medium, ion spray voltage of −4 500 V, temperature of 550°C, ion source gas of 1:60, and ion source gas of 2:60.

Based on the Novogene database, samples were detected using multiple reaction monitoring. The data files generated by UPLC-MS/MS were processed using SCIEX OS v1.4 to integrate and correct the peak. The main parameters were set as follows: minimum peak height, 500; signal/noise ratio, 5; and Gaussian smooth width, 1. The area of each peak represents the relative content of the corresponding substance. All central carbon substances were tested, and their categories are listed in Table S11.

### Molecular docking and molecular dynamics simulation

The protease sequence of LuxR solo was downloaded from GenBank (accession no: WP_003076066.1) in FASTA format. To build the 3D model of the LuxR solo protease, the target sequence information was submitted to the SWISS-MODEL server^80^ (http://swissmodel.expasy.org). Templates showing the highest quality were selected for model building. The predicted model output generated as a PDB file was downloaded for further analysis and visualized using SPDBV v4.10^81^. Structural coordinates of the potential protease activators were separated from the crystal structure of the *SdiA* protease in a complex with 3-oxo-C6-homoserine lactone (PDB ID: 482), available from the Protein Data Bank. Molecular docking simulations were used to explore the binding mode of the sulfonyl-containing compounds (Table S7) onto the 3D model of the LuxR solo protease using AUTODOCK tools v1.5.6 ^82^. Before docking, polar-H atoms were added to the LuxR solo model, and the macromolecule file was then saved in pdbqt format to be used for docking. The AutoGrid program generated ligand-centered maps with a grid dimension of 40 × 40 × 40. The Gridbox center was set to the x, y, and z coordinates −4.297, 17.693, and −27.541, respectively. Polar H charges of the Gasteiger type were assigned, non-polar-H atoms were merged with the carbons, and internal degrees of freedom and torsions were set. Default settings were used for all other parameters. The PyMol package^83^ was used to visualize the binding interactions between these ligands and the 3D LuxR solo protease.

### Western blotting to assess LuxR solo solubility

*LuxR solo* gene was PCR-amplified and digested with Xbal/Xhol, with the fragment then introduced into Xbal/Xhol-treated pET-22b(+) to generate pET-22b(+)-*LuxR*. The sequence-validated plasmid was used to transform *E. coli* BL21(DE3) cells for expression. A sterile culture tube containing NB broth supplemented with ampicillin was inoculated with a single colony of *E. coli* BL21(DE3) cells carrying pET-22b(+)-*LuxR*. Overnight cultures were inoculated into 100 mL of fresh medium and grown at 37°C/200 rpm to an OD_600_ of ∼0.6. Isopropyl-β-D-thiogalactoside (0.5 mM) was added to the cultures, which were subsequently divided into four 10-mL aliquots. Two aliquots received 75 μM SMX or SD, and two received an equivalent volume of dimethyl sulfoxide (0.1% vol/vol). After shaking at 20°C/180 rpm for 20 h, the cells were harvested by centrifugation (4 000 × *g*, 10 min, 4°C), resuspended in 2 mL of PBS buffer, and disrupted using an ultrasonic cell crusher (JY88-II). For protein solubility assessment, total cell lysates were separated into soluble and pellet fractions by ultracentrifugation at 4°C/45 000 × *g* for 30 min. Protein samples from the soluble and pellet fractions were fractionated by sodium dodecyl sulfate-polyacrylamide gel electrophoresis (SDS-PAGE) and western blotting.

### Identification of *lux-box*

To determine the presence of *lux-box* in specific promoters of FAO genes, upstream sequences were retrieved using RSAT tools^84^, and promoter regions were identified using BPROM^85^. Twenty base pairs of palindromic sequences in the promoters were then identified using the motif discovery tool of MEME^60^. Identified sequences were then aligned with known *lux-box* sequences.

### Purification of LuxR solo protein

Purification of LuxR solo was carried out at 4°C using AKTA pure 25 M1 (Cytiva). The SMX-induced *E. coli* BL21(DE3) pET-22b(+)-*LuxR* cell pellet was resuspended in lysis buffer (50 mM Tris, 300 mM NaCl, 0.1% Triton X-100, 0.2 mM phenylmethylsulfonyl fluoride, pH 8.0). Subsequently, the suspension was stirred for 30 min and sonicated on ice for 2 min in 15-s on/15-s off cycles at 30% power. The sonication cycle was repeated after the suspension rested on ice for 5 min, after which cell debris was pelleted via centrifugation (8 000 × *g*, 20 min, 4°C). The crude extract was then loaded onto a nickel metal affinity column (5 mL) equilibrated with 5 CV binding buffer (50 mM Tris, 300 mM NaCl, pH 8.0) and incubated for 1 h at room temperature. Finally, the protein was eluted with elution buffer (50 mM Tris, 300 mM NaCl, 200 mM imidazole, pH 8.0), and the effluent fraction was collected. His-LuxR was dialyzed into protein preservation buffer (50 mM Tris, 300 mM NaCl, 0.1% sarkosyl, 2 mM DTT, pH 8.0), concentrated after dialysis with PEG 20000, and filtered through a 0.45-μm membrane. The desired protein fractions were pooled, analyzed by SDS-PAGE and UV-vis spectroscopy, dispensed in 1 mL/tube, and stored at −80°C.

### EMSA

For the preparation of fluorescent FAM-labeled probes, the promoter region of the pccdA-GFP plasmid was PCR-amplified with Dpx DNA polymerase from the plasmid 3-ketoacyl-CoA thiolase (KAT) using the M13F/M13R primers listed in Table S9. The FAM-labeled probes were purified using the Wizard® SV Gel and PCR Clean-Up System (Promega, USA) and quantified with a NanoDrop 2000C (Thermo, USA). EMSA was performed in a 20-µL reaction volume containing 50 ng of probe and various proteins in a reaction buffer of 50 mM Tris-HCl [pH 8.0], 100 mM KCl, 2.5 mM MgCl_2_, 0.2 mM DTT, 2 μg of polydIdC, and 10% glycerol. After incubation for 30 min at room temperature, the reaction system was loaded into a 6% PAGE gel buffered with 0.5× Tri-boric acid. Gels were scanned using an ImageQuant LAS 4000 mini (GE Healthcare).

### Quantification and statistical analysis

All statistical analyses were performed in R. Analysis details can be found in figure legends and the result section. Figures were prepared using the basic R package and ggplot2. Significance (\**p* < 0.05) was determined using the Wilcox test.

## Data availability

- The raw RNA-seq data were deposited in the NCBI GEO Short Read Archive (SRA) under accession numbers PRJNA933284.
- Any additional information required to reanalyze the data reported in this paper is available from the lead contact upon request.

## Acknowledgements

This work was supported by the National Natural Science Foundation of China (52293442 and 52250056) and the National Key R&D Program of China (2021YFC3200603).

